# Can Transcranial Direct Current Stimulation Over the Dorsolateral Prefrontal Cortex Enhance Proprioception?

**DOI:** 10.1101/539510

**Authors:** Eric N. Beck, Sankirtana Shankar Narayanan, Rian McDermott, Alice G. Witney

## Abstract

**Introduction:** Proprioception (perception of one’s limb position) is critical for accurate and consistent movement, and is processed by the sensorimotor cortex. Increased prefrontal activity is associated with improved proprioception and motor performance. Anodal transcranial direct current stimulation (tDCS) of the left dorsolateral prefrontal cortex (DLPFC) has been found to increase activity of the sensorimotor cortex. Thus, this study aimed to investigate whether anodal tDCS of the DLPFC may enhance proprioception measured with a target task. It was hypothesized that tDCS over the left DLPFC would improve motor performance (error and variability) on a target task completed without vision.

**Design:** Single blind, within-participant, sham-controlled trial.

**Methods:** Fifteen healthy young adults (M:F=6:9, age=23.3 years) completed 18 trials of a computerized target task (manipulating a mouse) with their non-dominant upper-limb, with and without vision, before and after (pre/post assessment) 20-minutes of stimulation (anodal tDCS of the left DLPFC) and sham conditions. Averages and coefficient of variation (CV, variability between trials) of spatio-temporal parameters associated with the movement were measured. Stimulation/ sham sessions were counterbalanced (stimulation first session, n=8), with each session separated by one week. Repeated-measures ANOVA and pairwise comparisons (95% confidence intervals [CI]) were conducted.

**Results:** Regarding distance travelled CV, a significant interaction between condition and assessment (F(1,14)=5.09, p=0.041) demonstrated that variability was significantly less post-stimulation compared to pre (p=0.003). A significant interaction between assessment and vision (F(1,14)=30.08, p<0.001) regarding distance travelled CV showed that without vision, variability was significantly less at post compared to pre (p<0.001), and this decrease was found after the stimulation condition only (95% CI = Δ 7.4 +/− 1.6 [4.0 to 10.9]).

**Conclusion:** Since variability of distance travelled during the target task without vision was lower post-stimulation compared to pre, consistency of movement without vision, and therefore proprioception, may have been enhanced by anodal tDCS of the DLPFC. This improvement could be due to modulation of fronto-striatal-thalamic circuits. These findings may be the first step in developing tDCS methods as an effective adjunct therapy for dysfunctional proprioception in various disorders, such as Parkinson’s disease.

## 1. INTRODUCTION

Proprioception refers to the perception of one’s limb position in physical space (1–3). To perform novel tasks, humans utilize multiple sensory modalities to ensure accuracy of movement, such as visual, vestibular, and proprioceptive information, the most relied upon being vision (4). Reliance on each sensory modality is not static but rather dynamic, and the degree to which we utilize each sensory modality depends on the task performed (5–7). For example, when vision is removed (such as when one’s eyes are closed), reliance on proprioceptive feedback to guide movement becomes predominant (5). With effective use of proprioception, individuals can make precise movements with little variability, despite lack of vision (1). However, in cases of impaired processing of proprioception (such as in Parkinson’s disease), movement accuracy decreases and variability increases (3, 8–12), which could potentially lead to tripping, falling, and hospitalization (13–17). Interestingly, individuals with Parkinson’s disease are thought to compensate for postural deficits by increasing activation of prefrontal cortex activity (18). Thus, an understanding of the neurophysiological processes involved in proprioception and the importance of fronto-striato-thalamic pathways are vital to establish future therapies for individuals with impaired perception of limb position in physical space.

Proprioceptive input from peripheral receptors is transmitted via the dorsal column and spinocerebellar pathways to the thalamus and then onto the somatosensory cortex (19). Based on studies in Parkinson’s disease, proprioception relies heavily on sensorimotor loops through the basal ganglia (8, 9, 20, 21). Deficits in cerebro-basal ganglia circuitry may be compensated for via increased activity in frontal striatal pathways (18, 19, 22). An increase in alpha power observed via EEG in the prefrontal cortex has been associated with proprioceptive training (23) whilst increased activation in the sensorimotor cortex reflects improved performance at proprioceptive tasks. For instance, Iandolo and colleagues (2015) demonstrated that while healthy participants completed a proprioceptive matching task in which neural correlates were investigated via functional magnetic resonance imaging (fMRI), the sensorimotor cortex activation increased (24). Taken together, this previous work suggests that an increase in sensorimotor cortex activation could be enhanced, and that fronto-striato-thalamic pathways may be a means of mediating this enhancement. One way to potentially influence sensorimotor cortex functioning in a safe and non-invasive manner is with the use of transcranial direct current stimulation (tDCS).

Transcranial direct current stimulation refers to the application of electrical current through electrodes placed on specific regions of the scalp (25, 26). The current passes through the scalp and has been postulated to modulate membrane potential, thus leading to increased or decreased neuronal excitability (27, 28). Stimulation via tDCS may be applied with either a positive current (anodal tDCS) that facilitates neuronal excitability or a negative current (cathodal tDCS) (25) that inhibits neuronal excitability. Therefore, one might predict that anodal current could be utilized to modulate sensorimotor excitability, and thus influence proprioceptive processing. Various brain regions could be targeted with anodal stimulation, although as previously discussed, fronto-striato-thalamic pathways may be a means of mediating proprioceptive enhancement. One prominent area of interest in sensory modulation has recently been the left dorsolateral prefrontal cortex (DLPFC). The DLPFC is a critical region for cognitive and emotional processing, properties previously exploited with anodal tDCS to modulate working memory (29, 30), sustained attention (31), depression (32, 33), and various other processes in both healthy and patient populations (34). Importantly, manipulation of neuronal excitability in the DLPFC has been found to modulate sensory perception, such as pain, fibromyalgia (35, 36), and more recently, tinnitus (37). It should be noted that these diverse effects are not the result of separate networks, but rather interconnected circuitries, demonstrated recently by Deldar et al. (2018) whom concluded that anodal tDCS over the left DLPFC improved pain via enhanced working memory (38). The non-specific cognitive, emotional, and sensory modulatory effects of tDCS over the DLPFC can be attributed to both the poor localisation of tDCS and the modulation of interconnected networks, as opposed to only local cortical areas underlying the electrodes (34).

In a novel study, Stagg and colleagues (2013) demonstrated that twenty minutes of anodal stimulation over the left DLPFC resulted specifically in increased perfusion of the sensorimotor cortices bilaterally, indicating greater functional connectivity between the left DLPFC and the sensorimotor cortex (measured with magnetic resonance imaging) (39). Since increased activity of the sensorimotor cortex may indicate improvement to proprioceptive processing (40), anodal tDCS over the left DLPFC might be expected to result in improved proprioception in a group of healthy individuals. In addition to increased coupling between the DLPFC and the sensorimotor cortex, Stagg and colleagues (2013) found decreased functional connectivity between the left DLPFC and the thalamus after anodal tDCS over the left DLPFC. This decrease in coupling between the DLPFC and the thalamus has been suggested to be a key contributor underlying modulation of sensory thresholds (i.e. decreased perception of pain) after tDCS over the left DLPFC (35, 39, 41), and might be expected to hinder proprioception. However, increased activity in the thalamus in individuals with Parkinson’s disease with chronic deep brain stimulation has been shown to lead to proprioceptive deficits suggesting that this stimulation could result in thalamic, thalamocortical or corticothalamic connectivity alterations that impair proprioception (42). The disruption of sensory processing in individuals with Parkinson’s disease has been suggested to be due to a deficit in sensory gating which may explain why both increases and decreases in sensory thresholds appear to be observed in this cohort. Since anodal tDCS over the DLPFC is commonly used to modulate sensory thresholds, the question as to whether tDCS may enhance proprioception is a timely and important inquiry to gain further insight into mechanisms underlying sensory modulation.

Various methods have previously been utilized to assess proprioception and thus make inferences regarding proprioceptive processing (1). One common method is a joint position reproduction task (43–45). For this task, a participant’s limb begins in a neutral position that is subsequently moved, either passively or actively, to a target joint position. The participant’s limb is then returned to the neutral position in which the participant’s goal is to return his/her limb to the remembered target position, all of which is accomplished without the use of vision. Effective proprioception would result in minimal error between the target position and the performed position with minimal variability between trials (1). Similarly, targets may be utilized in which a participant is required to move his/her limb or an object to a target with and without the use of visual feedback, referred to as a visuomotor target task (10). Again, minimal error between the target position and performed position with minimal variability between trials is indicative of effective proprioception.

Therefore, the aim of the present study was to investigate the influence of anodal transcranial direct current stimulation applied over the left dorsolateral prefrontal cortex on proprioception in healthy adults, assessed with the use of a visuomotor target task. It was expected that if tDCS over the left DLPFC does increase activity of the sensorimotor cortex that is involved in the perception of proprioceptive information, than performance (error and variability) on a target task completed without vision might improve (i.e. decreased error and variability compared to baseline). In contrast, if tDCS over the left DLPFC decreases activity of the thalamus involved in projecting sensory information to various regions of the cortex and suppressing erroneous information, than performance on a target task completed without vision might be hindered (i.e. increased error and variability compared to baseline).

## 2. METHODS

### 2.1. Participants

Fifteen healthy young adults (M:F=6:9; Right handed:Left handed= 12:3; age=23.3 years) participated in the present single blind, within participant, sham-controlled study. Members of the Trinity College Dublin community were recruited by word-of-mouth. Participants were included if they were between the ages of 18 and 30 years, and had not been diagnosed with any medical condition. Participants were excluded if they had a history of epilepsy, fainting, syncope, head trauma, severe headaches, movement disorders, or neuropathy. The Faculty Research Ethics Committee of Trinity College Dublin approved this study. All participants were informed of the experimental protocol. Written consent was obtained according to the Declaration of Helsinki prior to testing.

### 2.2. Experimental Setup

Participants were asked to visit the lab on two separate occasions, one week apart, to complete a visuomotor target task (task description to follow) before and after (pre- / post-assessment) a stimulation or sham condition (stimulation and sham protocol description to follow). At the beginning of the first session, each participant completed a medical health questionnaire form to determine eligibility and to sign the informed consent. Furthermore, participants completed the Waterloo Handedness Questionnaire (47) to determine degree of upper-limb dominance. Participants additionally completed the Trail-Making Task (parts A and B) (48) and the digit span working memory task. Whether completion of the stimulation or sham condition took place on the first or second visit to the lab was randomized and counterbalanced (8 participants completed the stimulation condition on the first session). Each participant completed both stimulation and sham conditions. Participants began each session by completing baseline evaluation of proprioception with the visuomotor target task. Subsequently, participants received twenty minutes of either anodal tDCS stimulation over the left DLPFC, or twenty minutes of sham. After completion of either stimulation or sham, participants completed the visuomotor target task again.

### 2.3. Transcranial Direct Current Stimulation

Transcranial direct current stimulation (tDCS) is a safe, non-invasive neuromodulatory technique that delivers a low current to the scalp. Since the purpose of the study was to apply current over the left DLPFC in a similar fashion to that completed by Stagg and colleagues (2013), the tDCS method utilized in the current study followed a similar protocol. A NeuroConn DC stimulator (neuroConn GmbH, Germany) was utilized to apply current. The anodal electrode was positioned on the F3 position (using the 10/20 EEG system for positioning transcranial magnetic stimulation) while the cathodal electrode was placed on the contralateral supraorbital ridge (36, 39, 49). Each 5 × 7cm electrode transferred current to the scalp via a saline-soaked surface sponge (49). In the stimulation condition, participants received 20 minutes of 1mA current with fade-in/fade-out periods of 10 seconds while seated (39). In the sham condition, participants were subjected to an identical protocol as the stimulation condition. However, participants received only 30 seconds of 1mA current with fade-in/fade-out periods of 10 seconds, followed by 19minutes, 30seconds of no stimulation. Participants were blinded to whether they received stimulation or the sham condition. To provide an indication of current distribution through the brain using the electrode montage described above (i.e. anodal over F3 and cathodal over supraorbital ridge), HD-Explore Neurotargeting software (HD-Explore Version 2.1, Soterix Medical) was utilized.

### 2.4. Visuomotor Target Task to Assess Proprioception

To measure proprioception with a visuomotor target task, Matlab R2017b (The MathWorks, Natick, MA) and Psychtoolbox-3 programs were utilized. Matlab / Psychtoolbox code were used to generate three separate coloured squares situated in different locations of a monitor that served as the targets (in the upper right corner [blue], upper left corner [red], and upper middle [green]; Fig 1). Participants sat at a desk in a comfortable position in front of the computer monitor, which presented the three squares. Participants were asked to move a computer mouse that manipulated the cursor on the monitor using their non-dominant upper-limb throughout the task. Use of the non-dominant upper-limb was chosen to increase task difficulty so as to avoid ceiling effects. The goal of the task, with each trial, was to move the cursor on the monitor (with the mouse) from the starting point, situated at the bottom middle position, to the centre of one square. Before each trial, the investigator instructed the participant as to which square was the target of that trial, and whether they were able to use visual information. In trials where vision was permitted, participants completed the task by moving the mouse from the starting position to the middle of the designated square for that trial. Once the participant believed they were in the correct position, the trial was completed. In trials where vision was not permitted, participants were given 2 seconds to see the position of the square, then were asked to move a blindfold over their eyes with their dominant upper-limb, and subsequently move the mouse/cursor with their non-dominant upper-limb to the position they believed was the middle of the designated square. Once the cursor was in the position they believed was the middle of the square, the trial ended. Prior to starting the task, the participant completed five practice trials for each box to become familiarized with the task. Participants completed 3 trials for each square (left, middle, right), both with and without vision (vision vs. no vision), before and after (pre vs. post assessment) stimulation and sham conditions, for a total of 72 trials. Participants were not provided knowledge of results.

**Figure 1:**
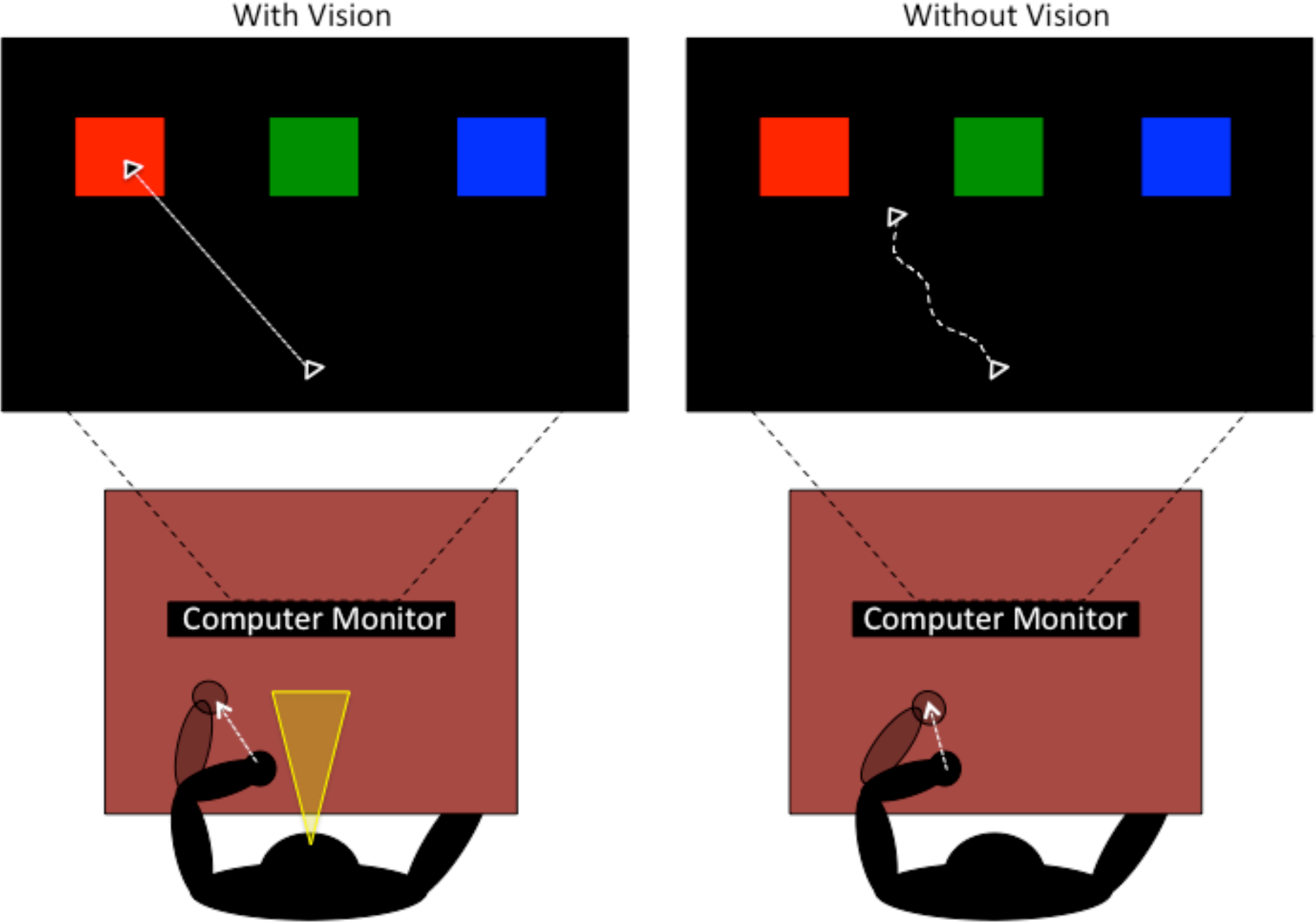
Visuomotor task setup. Participants sat at a desk in front of the computer monitor, which presented the three squares. Participants manipulated the cursor on the monitor from the starting point to the centre of one square, i.e. the target, with their non-dominant hand. Once the participant believed they were in the correct position, the trial was completed. In trials where vision was not permitted, participants were given 2 seconds to view the position of the square, then moved a blindfold over their eyes with their dominant hand, and subsequently moved the mouse/cursor to the position they believed was the middle of the designated square. Once the cursor was in the position they believed was the middle of the square, the trial ended.

### 2.5. Data and Statistical Analysis

The Matlab output provided x-y pixel coordinates (converted to millimeters [x/3.795]) of the mouse cursor collected at 60Hz. From this output, the following upper-limb movement parameters were calculated: i) spatial error of the trial endpoint compared to the middle of the designated square (millimeters [mm]); ii) movement time (seconds) comprising the total time elapsed from movement initiation to movement cessation; iii) distance travelled (mm) from movement initiation to movement cessation; iv) velocity (mm/second); v) x-y r-squared (spatial variability throughout the movement path where greater values indicate lower variability); vi) x-time R squared and y-time r-squared (spatial/temporal variability throughout the movement path). Within each parameter, data from each trial regarding each square target (left, middle and right) was collapsed to establish an average and coefficient of variation (CV = (standard deviation/mean) × 100) between trials.

To investigate the influence of anodal tDCS over the DLPFC on the average and variability (CV) of upper-limb movement (with respect to the target) spatial error, time, distance, velocity, and r-squared (x-y; x-time; and y-time), three-factor (condition [stimulation vs. sham] x assessment [pre vs. post] x vision [vision vs. no vision]) mixed repeated measures analysis of variance (ANOVA) were utilized. To determine where significant differences were with respect to main effects and interactions, a Fisher’s Least Significant Difference (LSD) post hoc analysis was used. Additionally, pairwise comparisons within (differences between pre and post assessment; differences between vision and no vision) and between the stimulation and sham means were conducted (95% Confidence Intervals [CI]) with regard to each upper limb movement parameter. All results were analyzed using StatSoft STATISTICA 8.0.550 (StatSoft Inc, Tulsa, Oklahoma) and the level of significant difference was set to p=0.05.

## 3. RESULTS

Participant demographics are presented in table 1 with handedness score, trail making parts A, B, and B-A times, and digit span working memory scores. Please see supplementary material for a table presenting all findings with respect to the proprioception task movement parameters, including significant main effects and interactions with partial-eta^2^.

**Table 1:** Participant Demographics

| Participant | Sex | Age | Waterloo<br>Handedness<br>Questionnaire | H | TMT<br>A | TMT<br>B | TMT<br>B-A | Digit<br>Span | COUNTER<br>BALANCE<br>START |
| --- | --- | --- | --- | --- | --- | --- | --- | --- | --- |
| 1 | M | 24 | 60 | R | 24.92 | 35.51 | 10.59 | 18 | STIM |
| 2 | F | 23 | -60 | L | 18.75 | 33.89 | 15.14 | 15 | STIM |
| 3 | M | 20 | 30 | R | 16.38 | 22.96 | 6.58 | 14 | SHAM |
| 4 | F | 24 | -16 | L | 15.58 | 37.08 | 21.5 | 25 | SHAM |
| 5 | F | 20 | 47 | R | 11.71 | 26.63 | 14.92 | 12 | SHAM |
| 6 | M | 24 | 39 | R | 15.71 | 26.48 | 10.77 | 18 | STIM |
| 7 | F | 21 | 64 | R | 14.28 | 33.08 | 18.8 | 16 | STIM |
| 8 | M | 27 | 20 | R | 16.95 | 49.38 | 32.43 | 10 | STIM |
| 9 | F | 26 | 48 | R | 12.35 | 21.8 | 9.45 | 19 | STIM |
| 10 | F | 26 | 47 | R | 15.01 | 56.11 | 41.1 | 17 | STIM |
| 11 | F | 20 | 62 | R | 14.71 | 31.95 | 17.24 | 19 | STIM |
| 12 | M | 25 | 56 | R | 11.38 | 30.97 | 19.59 | 22 | SHAM |
| 13 | F | 20 | 44 | R | 15.1 | 27.48 | 12.38 | 26 | SHAM |
| 14 | F | 20 | -56 | L | 22.46 | 39.37 | 16.91 | 18 | SHAM |
| 15 | M | 29 | 52 | R | 11.62 | 30.2 | 18.58 | 20 | SHAM |

H=Handedness; TMT A, B, and B-A = Trail-Making Task Part A, B, and B-A

With respect to modeled current distribution through the brain with HD-Explore Neurotargeting software, estimated current flow and magnitude for the anodal-F3 and cathodal-supraorbital montage is illustrated in figure 2. By characterizing the estimated current flow employed by the montage of the present study, potential brain regions affected by stimulation may be highlighted. Figure 2 demonstrates that anodal tDCS over the left DLPFC and cathodal tDCS over the right supraorbital ridge was expected to produce current flow that projected into the frontal lobe (0.15-0.2 V/m) to the anterior cingulate cortex (0.15-0.2 V/m), genu of the corpus callosum and varying subcortical regions (0.15-0.2 V/m), and various areas of the sensorimotor cortex (0.075-0.185 V/m).

**Figure 2:**
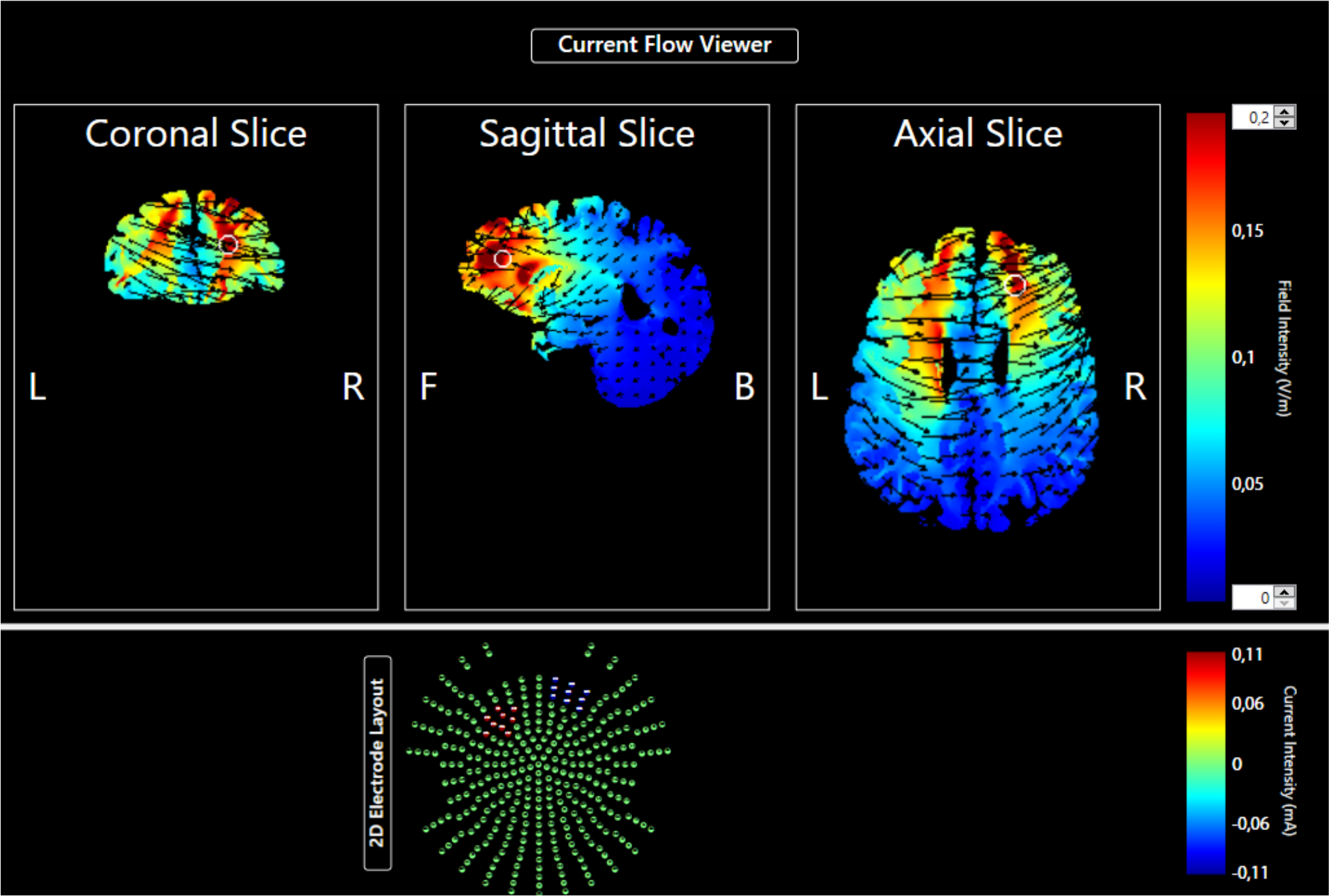
tDCS electrode current flow modelling for the current study. 2D Electrode Layout (bottom illustration): Electrode in Red (Anode) corresponds to position F3. Electrode in blue (cathode) corresponds to positioning over the supraorbital ridge. Current Flow Viewer (Top Illustrations): Characteristic flow of current, viewed through 2D coronal, sagittal and transverse (axial) slices. Current magnitude is illustrated by field intensity (V/m) on a continuum wherein blue signifies low intensity of current flow to red signifying higher current flow.

With respect to all movement parameters measured during the visuomotor target task, multiple significant main effects and interactions were uncovered. Firstly, to demonstrate that blindfolding participants did remove visual input and influenced an increased reliance on proprioception to guide movement, main effects of vision were found with respect to average spatial error (F(1,14)=799.4, p<0.001), velocity CV (F(1,14)=36.24, p<0.001), average x-y r-squared (F(1,14)=27.21, p<0.001), x-y r-squared CV (F(1,14)=12.09, p=0.004), average x-time r-squared (F(1,14)=23.29, p<0.001), x-time r-squared CV (F(1,14)=13.39, p=0.003), average y-time r-squared (F(1,14)=5.66, p=0.032), and y-time r-squared CV (F(1,14)=6.48, p=0.023). Specifically, throughout the visuomotor target task, participants demonstrated significantly greater average spatial error, x-y r-squared, x-time r-squared, and y-time r-squared when the task was performed without vision compared to with vision. Additionally, when performing the visuomotor target task with vision, variability between trials was significantly greater with respect to velocity, x-y r-squared, x-time r-squared, and y-time r-squared compared to no vision.

Main effects of assessment (pre vs. post) were found with respect to movement time (F(1,14)=5.31, p=0.037) and velocity (F(1,14)=5.54, p=0.034), such that at post assessment, participants performed the visuomotor target task with a significantly shorter movement time and greater velocity. It should be noted that pairwise comparisons (95% CI) between pre and post assessment demonstrated that with vision in the sham condition, movement time decreased from pre assessment to post (95% CI = 0.3 +/− 1.4 [0.1 to 0.6]), although not in the stimulation condition. Additionally, with and without vision in the sham condition, velocity increased from pre assessment to post (95% CI vision = −10.9 +/− 4.8 [−21.1 to −0.7]; no vision = −12.1 +/− 4.8 [−22.3 to −1.8]).

A significant interaction between condition, assessment, and vision was found with respect to variability (CV) of spatial error between trials (F(1,14)=8.75, p=0.011; Fig 3), and Fisher’s post hoc uncovered a number of significant differences. Firstly, within the stimulation condition, at post assessment only, spatial error variability between trials was significantly greater when participants had the use of vision compared to when blindfolded (no vision) (p=0.027). Secondly, within the sham condition, at pre assessment, spatial error variability between trials was significantly greater when participants had the use of vision compared to when blindfolded (p=0.002). Moreover, spatial error variability with vision within the sham condition significantly decreased from pre assessment to post (p=0.02). Finally, spatial error variability with vision was significantly greater at pre assessment of the sham condition compared to pre assessment of the stimulation condition (p=0.013).

**Figure 3:**
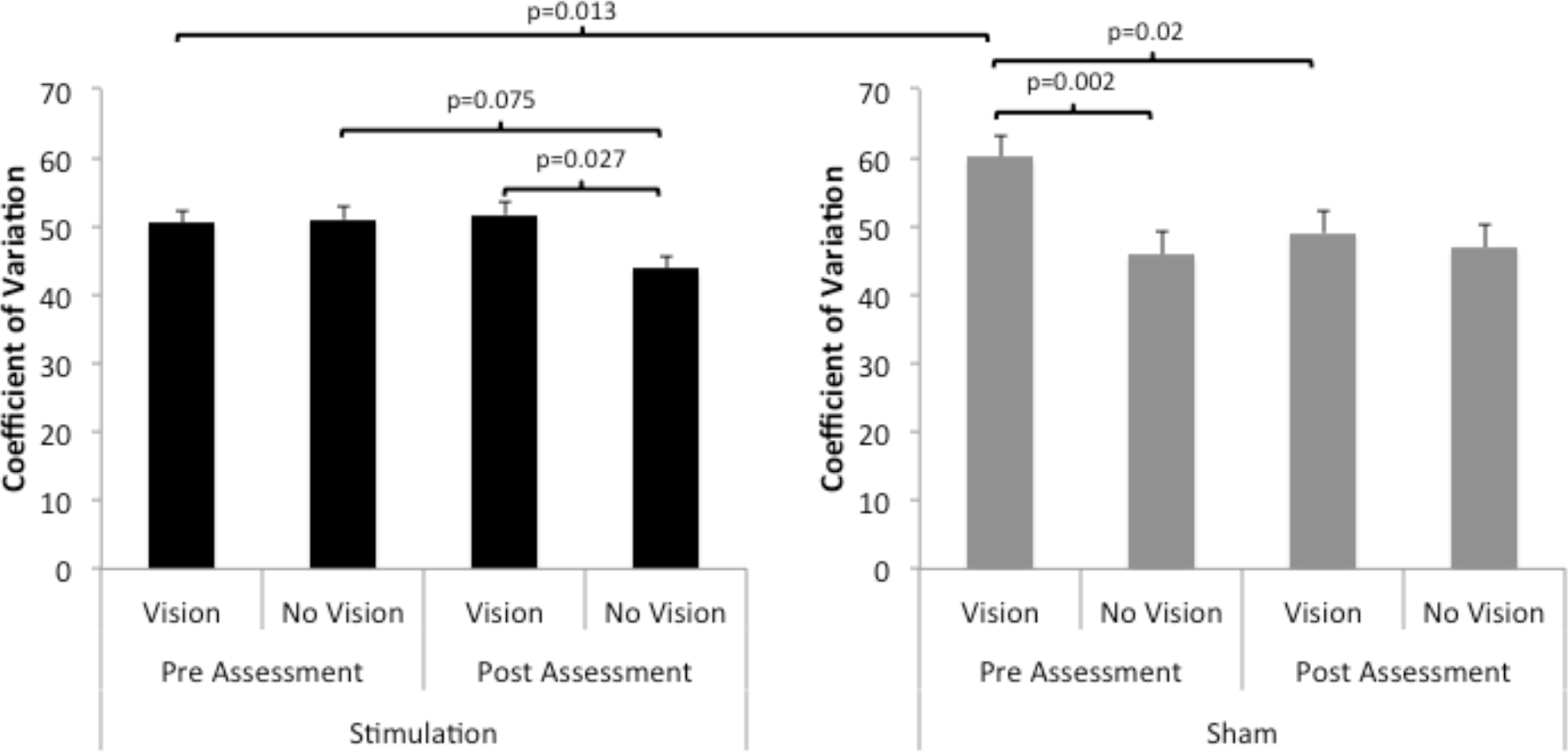
Significant interaction between condition (stimulation vs. sham), assessment (pre vs. post assessment), and vision (vision vs. no vision) with respect to the **spatial error coefficient of variation**.

Interestingly, regarding the variability of distance travelled (CV), significant interactions between condition and assessment (F(1,14)=5.09, p=0.041), and between assessment and vision (F(1,14)=30.08, p<0.001) were found (Fig 4). Fisher’s post hoc revealed three significant findings. Firstly, distance travelled variability between trials was significantly lower after stimulation (post) compared to before stimulation (pre) (p=0.003). Secondly, regardless of the condition or assessment, participants demonstrated significantly greater distance travelled variability without vision compared to with vision (p<0.001). Thirdly, when participants performed the task without vision, variability of distance travelled was significantly lower at post assessment compared to pre (p<0.001). Notably, pairwise comparisons demonstrated that the variability of distance travelled without vision decreased from pre assessment to post in the stimulation condition (95% CI = 7.4 +/− 1.6 [4.0 to 10.9]), and not the sham (95% CI = 1.2 +/− 1.6 [−2.2 to 4.7]). Variability of velocity between trials without vision also decreased from pre assessment to post within the stimulation condition (95% CI = 7.1 +/− 3.3 [0.1 to 14.1]), and not the sham (95% CI = 3.6 +/− 3.3 [−3.4 to 10.6]).

**Figure 4:**
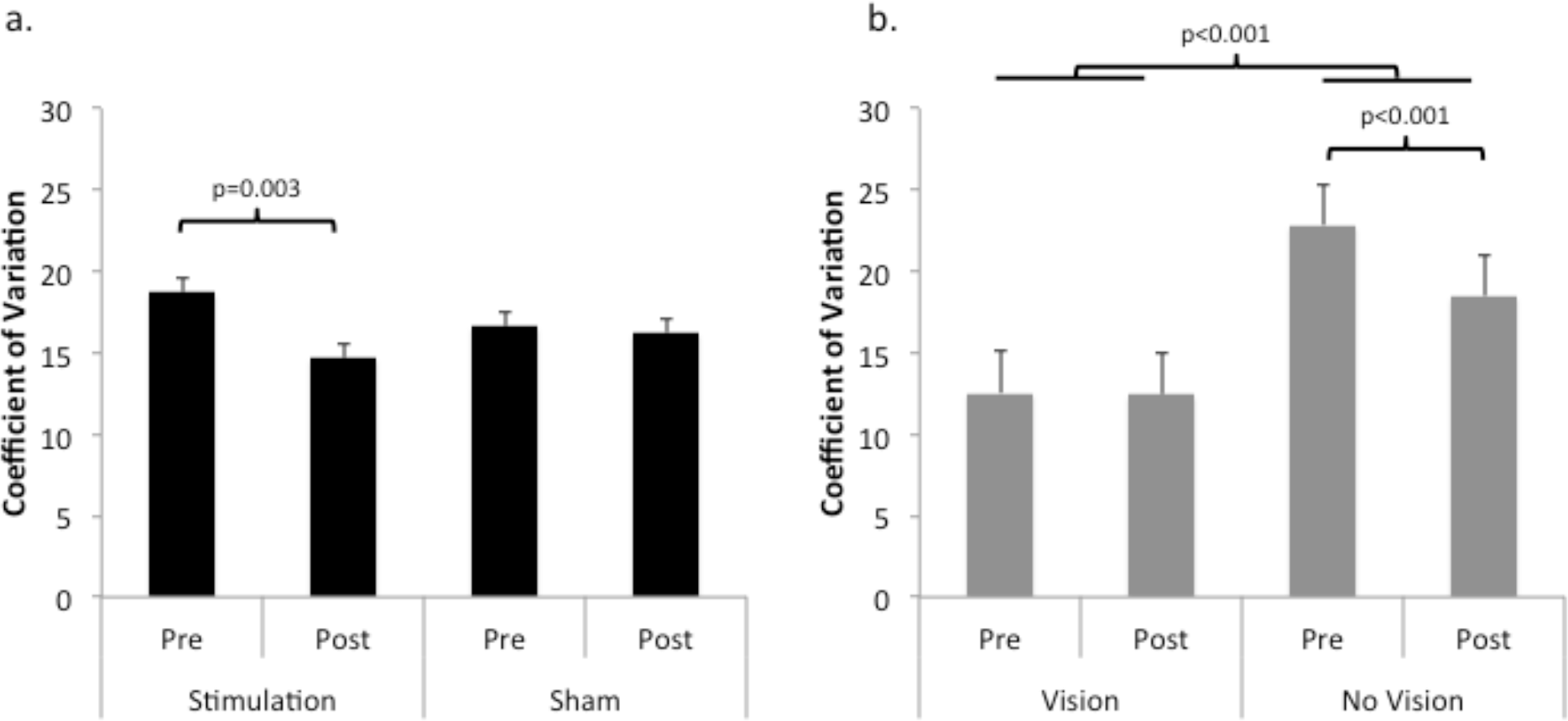
**Graph a:** Significant interaction between condition (stimulation vs. sham) and assessment (pre vs. post assessment) with respect to the **distance travelled coefficient of variation. Graph b:**Significant interaction between condition and assessment (pre vs. post assessment) and vision (vision vs. no vision) with respect to the **distance travelled coefficient of variation.**

Finally, a significant interaction was found between condition and assessment in regards to x-y r-squared variability between trials (F(1,14)=8.57, p=0.011; Fig 5). Fishers post hoc uncovered that x-y r-squared CV was significantly greater at post assessment compared to pre within the stimulation condition (p=0.004). Furthermore, x-y r-squared variability was significantly greater at post assessment of the stimulation condition compared to post assessment of the sham condition (p=0.017). Interestingly, pairwise comparisons demonstrated that the increase in x-y r-squared variability, from pre assessment to post, in the stimulation condition was only found when participants completed the task with vision (95% CI = −12.8 +/− 5.9 [−25.5 to −0.2]), and not without vision (95% CI = −6.3 +/− 5.9 [−19.0 to 6.3]).

**Figure 5:**
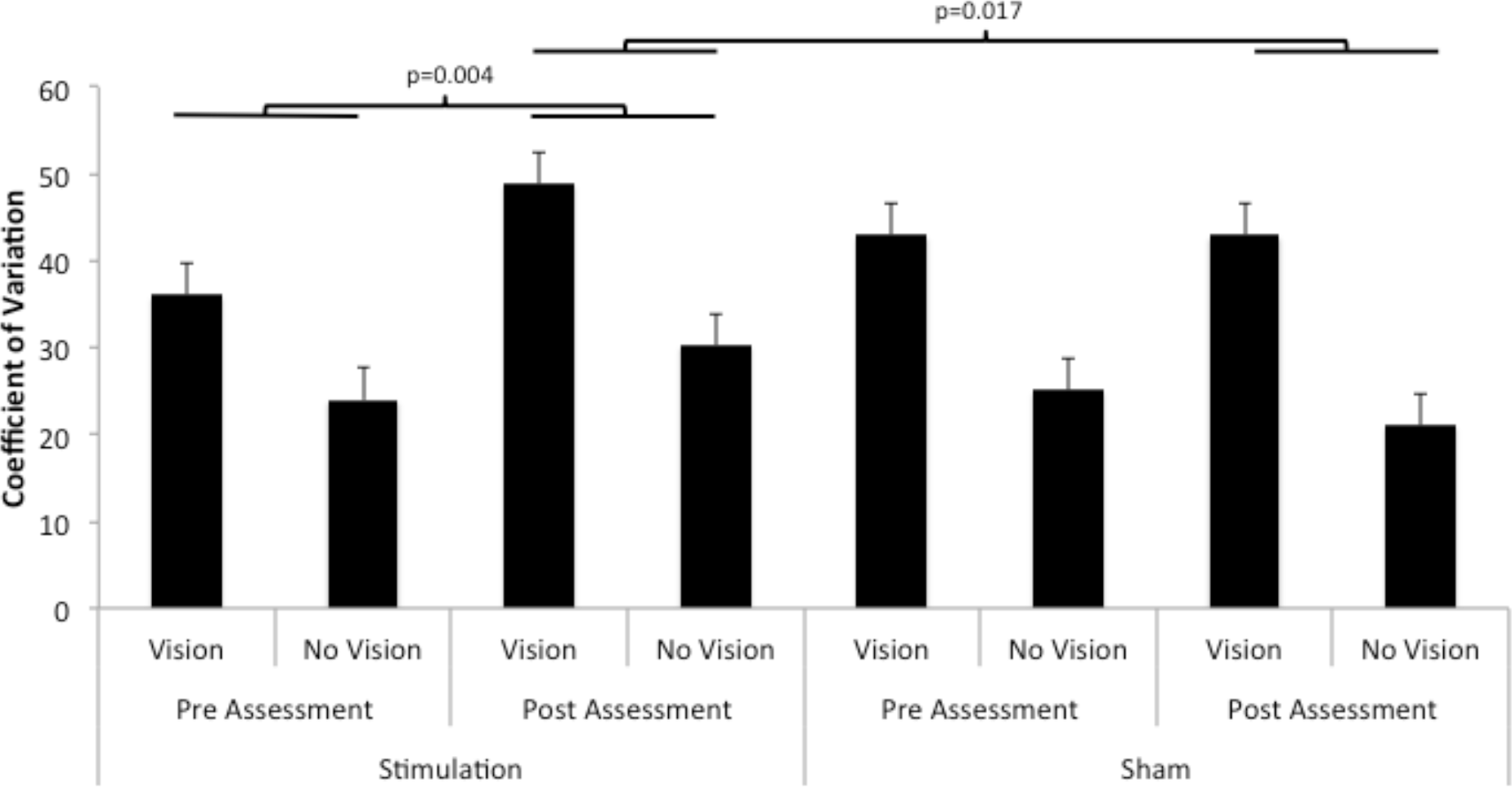
Significant interaction between condition (stimulation vs. sham) and assessment (pre vs. post assessment) with respect to the **X, Y R-Squared coefficient of variation.**

## 4. DISCUSSION

To our knowledge, this was the first study to investigate the influence of anodal transcranial direct current stimulation applied over the left dorsolateral prefrontal cortex on proprioception in healthy adults. It was hypothesized that if anodal tDCS over the left DLPFC does increase activity of the sensorimotor cortex that is involved in the perception of proprioceptive information, movement error and variability on a visuomotor target task performed without vision might improve. Interestingly, participants completed the visuomotor target task with significantly lower distance travelled variability after anodal tDCS compared to before (Fig 4a). When participants performed the task without vision, variability of distance travelled was significantly lower at post compared to pre (Fig 4b), and this decreased variability was found specifically after stimulation, but not sham. Therefore, after 20 minutes of anodal tDCS of the left DLPFC, variability of distance travelled during the visuomotor target task without vision had decreased, indicating that consistency of movement without vision improved. Typically, as proprioception is enhanced or becomes more relied upon, conscious adjustments of movement decrease and consistency of movement performance increases (50–54). Since this decrease in variability was not found after the sham condition, it is unlikely that the improved consistency was the result of performing the task a second time. In summary, consistency of upper-limb movement improved after anodal tDCS of the left DLPFC and this may be an indication of enhanced proprioception. The significant findings with regards to the other movement parameters provide further support to these inferences.

Before anodal tDCS of the DLPFC, spatial error variability during the visuomotor target task was similar between vision and no vision parameters. After anodal tDCS, spatial error variability was significantly lower when participants performed the target task without vision compared to with vision (Fig 3). Furthermore, from pre stimulation to post, without vision, spatial error variability anecdotally decreased, although this was only indicated by a trend (p=0.075). The decrease in spatial error variability may support the previous findings (distance travelled variability decrease after tDCS without vision) that indicate improved consistency between trials, and therefore enhanced proprioception after anodal tDCS of the left DLPFC that was not found after the sham condition. However, interpretations of these findings cannot be strongly made since spatial error variability with vision before the sham condition, was significantly greater than spatial error variability with vision before the stimulation condition. An explanation for this finding cannot be made since data collection was performed in an identical fashion prior to both stimulation and sham conditions and the experimental design was counterbalanced. However, since this aberrant finding was only found with the use of vision and not when participants performed the target task while blindfolded, spatial error variability, and therefore proprioception, may have improved after anodal tDCS of the left DLPFC and not after the sham.

The final finding of interest was a significant interaction between condition and assessment with respect to x-y r-squared variability between trials. The x-y r-squared is a quantitative measure of the spatial variability throughout the movement path where greater values indicate lower variability, and therefore fewer adjustments throughout the route from the starting point to the target. The coefficient of variation of the x-y r-squared is therefore a measure of variability between trials with respect to the amount participants made adjustments throughout each trial. Figure 5 demonstrates that from pre stimulation to post, x-y r-squared variability significantly increased, indicating that the amount of adjustments made throughout each trial was more variable after anodal tDCS compared to before. This finding might suggest that after stimulation, participants attended more to their movements throughout each trial, which resulted in a greater degree of variability between trials and may indicate a decrement in proprioception, contradicting our previous findings. However, pairwise comparisons demonstrated that the increase in x-y r-squared variability, from pre stimulation to post, was only found when participants completed the task with vision. This might suggest that the significant increase in variability from pre stimulation to post was driven by task performance with vision and not without. Nevertheless, these findings with respect to distance travelled variability, spatial error variability and x-y r-squared variability allow for various inferences.

Stagg and colleagues (2013) demonstrated that anodal tDCS over the left DLPFC resulted in increased activity of the sensorimotor cortex and decreased activity of the thalamus. One may have hypothesized that increased activity of the sensorimotor cortex might enhance proprioception, and decreased functional connectivity between the DLPFC and the thalamus (35, 39, 41) might hinder proprioception. Our results demonstrated that anodal tDCS of the left DLPFC resulted in decreased variability with respect to the end-point of each trial (i.e. decreased distance travelled CV and spatial error CV without vision after stimulation compared to before). Therefore, the modulation of proprioceptive processing in this study may have been enhanced by increased sensorimotor cortical activity induced by anodal tDCS over the DLPFC. Previous work has demonstrated that anodal tDCS, which facilitates neuronal excitability, significantly decreases concentrations of the inhibitory neurotransmitter gamma-amino butyric acid in local cortical areas underlying the anodal electrode (55). Down-regulated inhibition of the DLPFC may underlie various changes in networks associated with the DLPFC that yielded improved proprioception, such as enhanced sensorimotor activity (39, 55).

On the other hand, greater variability between trials with respect to the movement performed in reaching the end-point (i.e. increased x-y r-squared variability from pre assessment to post) might suggest that proprioceptive processing was hindered, supporting the notion that stimulation of the left DLPFC decreased activity of the thalamus projecting sensory information to various regions of the cortex, allowing irrelevant information to propagate through the thalamus that might interfere with proprioceptive processing (46). Although, another explanation for the increased variability with respect to x-y r-squared may be that stimulation of the left DLPFC influenced participants’ focus of attention. Previous work has demonstrated that the left DLPFC is recruited when individuals pay attention to performance of pre learned tasks (50). Moreover, anodal tDCS over the left DLPFC has been found to modulate various cognitive processes, such as working memory (29, 30) and sustained attention (31). Therefore, by stimulating the DLPFC, focus of attention towards enhanced online proprioceptive feedback during movement may have been increased, resulting in variability between trials. Alternatively, if proprioception became enhanced, participants may have gained improved ability to process that proprioceptive feedback online during movements, allowing them to make more fine tune adjustments (increased x-y r-squared variability) to achieve lower spatial and distance travelled variability.

To date, no study has directly aimed to enhance proprioception with the use of tDCS. However, multiple studies have investigated the influence of tDCS on postural control, a dynamic process in which integration of proprioception with visual and vestibular information is imperative for effective balance. Craig and Doumas (2017) recently demonstrated that anodal tDCS applied over the primary motor cortex or the cerebellum in young and older healthy adults did not improve postural control (56). Similarly, anodal tDCS over the primary motor cortex was not found to facilitate learning of a dynamic balance task, a phenomenon one might expect if tDCS enhanced sensorimotor cortical activity (57). In a very relevant contrast to these two null findings, Zhou and colleagues (2015) demonstrated that tDCS applied over the left DLPFC improved postural control while participants performed a balance task in single and dual-task (serial subtraction) conditions (58). These findings by Zhou et al. (2015) are critical in establishing the importance of anodal tDCS over the DLPFC, and not other sites such as the cerebellum or primary motor cortex, in order to improve proprioception.

Performance of a motor and cognitive task simultaneously is an effective method to determine the degree to which the motor task requires attention for successful performance (59). Since Zhou et al. (2015) found that postural control (motor task) while counting backwards by three’s (cognitive task) improved after anodal tDCS of the left DLPFC, this might indicate that postural control required less cognitive demand after stimulation (60, 61). Meanwhile, Stagg and colleagues (2013) demonstrated that anodal tDCS of the left DLPFC increased activity of the sensorimotor cortex (39). Thus, the improved postural control while dual-tasking found by Zhou et al. (2015) may have been the result of increased sensorimotor activity that improved proprioceptive processing, decreasing the requirement for attention to control balance. Importantly, these findings by Zhou and colleagues (2015) amalgamated with the findings by Stagg et al. (2013) further support that the improved consistency of upper-limb movement after anodal tDCS of the left DLPFC found in the present study may be indicative of enhanced proprioception.

### 4.1 Future Directions

The findings from the present study may be the first step in developing tDCS methods as an effective adjunct therapy for dysfunctional proprioception in various disorders, such as Parkinson’s disease (3, 8–12), wherein progressive neurodegeneration of dopamine producing cells of the basal ganglia takes place. Impaired processing of proprioception results in increased variability of walking (62–65), which leads to falling (15, 66), injury (16, 67), and even hospitalization (17). Dopaminergic replacement medications have been found to effectively manage motor symptoms in individuals with Parkinson’s disease, but are ineffective for the alleviation of increased walking variability (68, 69). Non-invasive neurostimulation of the prefrontal cortex has previously been demonstrated to modulate dopamine release in sub-cortical areas (70, 71). Previous research has indicated that prefrontal neurostimulation leads to improvements in working memory (72) and depression (73) in individuals with Parkinson’s disease, but some indication of a trend with regards to motor symptoms (74). These studies focused on clinical assessments of motor function and simple reaction time tasks rather than an assessment of changes in proprioception. Therefore, tDCS over the left DLPFC could in future studies be assessed as an effective adjunct therapy used to ameliorate proprioception deficits in individuals with Parkinson’s disease.

## Acknowledgements

We would like to thank Hannah Ashe for her assistance in setting up this study. We would also like to thank all of the individuals who took the time to participate.

